# Snow Microbiome Functional Analyses Reveal Novel Microbial Metabolism of Complex Organic Compounds

**DOI:** 10.1101/2020.02.07.938555

**Authors:** Chengsheng Zhu, Maximilian Miller, Nicholas Lusskin, Benoît Bergk Pinto, Lorrie Maccario, Max Haggblom, Timothy Vogel, Catherine Larose, Yana Bromberg

## Abstract

Microbes active in extremely cold environments are not as well explored as those of other extreme environments. Studies have revealed a substantial microbial diversity and identified cold-specific microbiome molecular functions. We analyzed the metagenomes and metatranscriptomes of twenty snow samples collected during early and late spring in Svalbard, Norway using our computational read-based microbiome function annotation tool, mi-faser. Our results revealed a more diverse microbiome functional capacity and activity in the early compared to in the late spring samples. The dissimilarity between the metagenomes and metatranscriptomes of the same samples was also significantly higher in the early spring. These findings suggest that early spring samples may contain a larger fraction of DNA of dormant organisms, while late spring samples reflect a new community that is metabolically active. We additionally showed that the abundance of the sequencing reads mapping to the fatty acid synthesis-related microbial pathways was significantly positively correlated with organic acid levels, in both our late spring metagenomes and metatranscriptomes. Moreover, the geraniol degradation pathway and the styrene degradation pathway read abundances correlated and inversely correlated, respectively, with the organic acid levels. These results suggest a possible nutrient switch. Our study thus highlights the activity of microbial degradation pathways of complex organic compounds previously unreported at low temperatures.

## 1. Introduction

Abiotic parameters, such as temperature, pH, and pressure create stress on microorganisms, especially in extreme environments [1]. The cryosphere, an extreme cold environment, covers a large portion of Earth’s surface. Over 14% of the world’s biosphere is located at the planetary poles, while 90% (by volume) of the ocean is colder than 5°C [2]. Taxonomic surveys based on 16S rRNA gene sequencing have described significant microbial diversity in glacial ice [3–5], cryoconite [6, 7], sea ice [8], and polar and alpine snow [9–13]. Bacteria seem to be ubiquitous in snow and belong to numerous taxa such as *Proteobacteria* (*Alpha*-, *Beta*- and *Gamma*-), the *Cytophaga-Flexibacter-Bacteriodes* group, *Actinobacteria*, and *Cyanobacteria [10, 12, 14, 15],* although the results can vary based on season, sampling location and analysis methods. For example, the diversity of organisms in snow from the Canadian high Arctic ice sheet was 20 times lower than that measured in Tibetan plateau snow *[12, 16]*. A variety of approaches, such as cultivation, ribosomal profiling, and stable isotope probing, have been used to detect and measure microbial activity at subzero temperatures in permafrost soils (for review, see Nikrad et al. 2016 [17]). These offer insights into the microbial interactions with the soil environment in the cold. Considerably less is known about the functional capacity of microorganisms in the snow. One pioneering metagenomic study identified microbiome functional capacity correlating with the chemical parameters (e.g. mercury concentration) in Arctic spring snow samples [18]. If organisms are active at below zero temperatures in the snow, then they are likely involved in a range of processes involving organic matter, which could impact atmospheric and biogeochemical cycles [19]. One poorly constrained source and potential modifier of organic compounds is biological activity in snow [20].

Large scale metagenome sequencing drastically increased the publicly available metagenomic data from high profile projects such as Terragenome [21], the Global Ocean Sampling Expedition [22], the Human Microbiome Project [23] and Tara Oceans [24]. Studies have been carried out in a wide variety of environments including the human gut [25–27], groundwater [28], acid mine drainage [29], beach sand [30], etc., and identified potential diagnostic, therapeutic, or bioremediation targets. With ample data, comparative analysis of meta-genomes/transcriptomes under different conditions highlights the key microbial members and functions that result from and/or contribute to niche differences [31, 32]. Comparative meta-genomes/transcriptomes analysis have not been commonly applied to cold environment samples. Such analysis should help elucidate microbial mechanisms of survival and adaptation at low temperatures.

Bergk-Pinto et al. studied the microbial ecology in twenty snow samples collected during early and late spring (mid-April to mid-June, 2011, Svalbard, Norway) [33]. Using a combined method of marker genes and network analysis, Bergk-Pinto et al. revealed that from early to late spring, the microbial community in the snow shifted from cooperation to competition, accompanied by enrichment of antibiotic resistant genes [33]. Here, we investigated microbial metabolism of organic compounds at low temperature and further analyzed these metagenomic and metatranscriptomic datasets using mi-faser **[27]**. This new bioinformatic tool allows functional annotation of sequencing reads based on experimentally verified microbial enzymes at high accuracy (>90%). Our results revealed significantly lower metagenome-to-metatranscriptome similarity in the early spring than in the late spring samples. We also found that in the late spring samples the abundance of sequencing reads mapping to the components of the fatty acid synthesis-related microbial pathways significantly correlated with organic acid levels. Both of these findings are consistent with a period of microbial community growth, in line with the previous finding of the switch from microbial cooperation to competition [33]. We further observed that the rise in organic acid levels correlated with the appearance of the geraniol degradation pathway and disappearance of the styrene degradation pathway. This finding might represent a change in nutrient conditions during the community growth process. To summarize, here we observed the presence of microbial functionality necessary for degradation of complex organic compounds in both metagenomes and metatranscriptomes of the late spring snow samples. This is the first time this functionality has been reported to be under *active expression* at temperatures below 0°C.

## 2. Materials and Methods

### 2.1. Data collection and preprocessing

We obtained the metagenomic, metatranscriptomic, and chemistry data for twenty snow samples from the ftp website from the Environmental Microbial Genomic Group (ftp://ftp-adn.ec-lyon.fr/Snow_organic_acids_bacterial_interactions/metagenomes_and_metatranscriptomes_svalbard2011/). The technical details of the sampling and sequencing process were described in a previous study [33]. Briefly, snow was collected over two months during mid-April to mid-June at Ny Ålesund in the Spitsbergen Island of Svalbard, Norway (78°56’N, 11°52’E). Surface snow layers were collected into sterile bags using a sterilized shovel from a 50 m2 perimeter with restricted access to reduce contamination from human sources. Details on sampling conditions, sample site and chemical analyses can be found in Bergk-Pinto et al. [33].

The sequencing data were processed using Mothur [34] for quality filtering using the settings in Schloss et al. [35]. Base overrepresentation was controlled using FastQC [36] and Usearch [37] was used to identify and remove remained adaptors.

### 2.2. Analysis

The post-quality-control samples were submitted to mi-faser web service **[27, 38]** for functional annotation. For each sample, mi-faser returns a read abundance table of enzyme functionality detected in the sample, the EC-profile (EC stands for Enzyme Commission **[39]**). For each known functional pathway **[40]**, we divided the sum of all reads mapping to the enzyme members of the pathway by the number of these enzyme members to create an entry in the pathway-profile of the sample. The NMDS diagrams were generated with the (enzyme and pathway) profiles of samples assigned to four groups, early_DNA (early spring metagenomes), early_RNA (early spring metatranscriptomes), late_DNA (late spring metagenomes), and late_RNA (late spring metatranscriptomes). The Euclidean distances between the same-sample DNA and RNA NMDS points were calculated and compared across groups. Note that our computational method (mi-faser followed by NMDS analysis) successfully revealed microbial functional diversity in various environment samples, such as beach sand and human gut **[27]**. The reliability of mi-faser annotation is from 1) its function mapping algorithm that is specifically trained for short reads, and 2) its manually curated reference database that contain only protein sequences with experimentally verified function **[27]**. Organic acid levels were standardized with their according total concentration in all samples. The Pearson correlation coefficients, as well as the significance of correlations, were calculated by the R function cor.test [41].

## 3. Results and Discussion

### 3.1. Early-to-late-spring dissimilarity and metagenome-to-transcriptome divergence highlight community activity in late spring samples

While the metagenome reflects the overall potential function of a microbial community, metatranscriptomic analyses are based on genes that are transcribed and thus provide more information on the active fraction of these functions. Using mi-faser (microbiome functional annotation of sequencing reads), our read based microbiome function annotation tool, to analyse the metagenomes and metatranscriptomes of early and late spring polar snow samples, we observed that (1) the early spring samples were more diverse (measured as the Euclidean distance between entries on the NMDS plot; Methods) in both potential and active microbial functionality than the late spring samples (EC-profiles sample distance: early spring = 4.8±2.3, late spring = 0.4±0.3, Figure 1A, SOM Figure 1A,B; pathway-profiles sample distance: early spring = 1.4±0.9, late spring = 0.1±0.1, Figure 1B, SOM Figure 1C,D) and that (2) meta-genome-to-transcriptome similarity of the same sample (measured as the Euclidean distance between entries on the NMDS plot; Methods) is was significantly lower in early than in late spring (in both comparisons of the EC-profiles, t-test p-value <0.001, Figure 2A and the pathway-profiles, p-value=0.025, Figure 2B). Note that for all comparisons ~29% ECs (195 of 683; SOM Table 1) in our data could not be mapped to known KEGG pathways.

**Figure 1.**
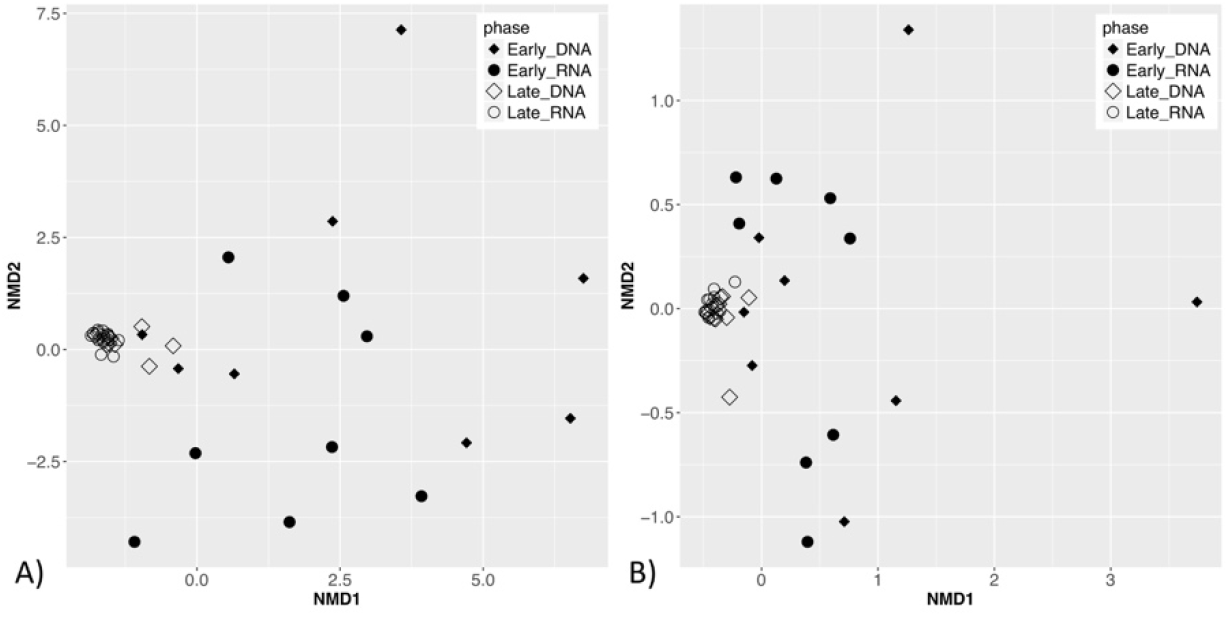
NMDS diagrams suggest higher microbial functional beta-diversity in the early spring samples than in the late spring ones. The diagrams represent A) EC-profiles. The average Euclidean distance is 4.8±2.3 between early spring samples, and 0.4±0.3 between late spring samples; B) pathway-profiles. The average Euclidean distance is 1.4±0.9 between early spring samples, and 0.1±0.1 between late spring samples.

**Figure 2.**
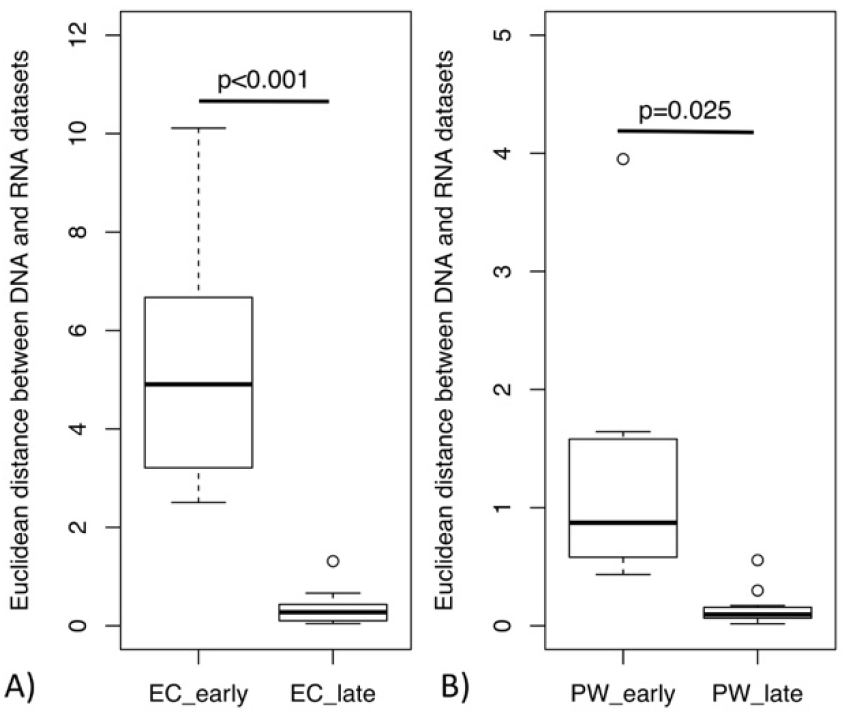
The metagenome-metatranscriptome distance (dissimilarity) in the early spring samples is significantly higher than in the late spring samples. The boxplots were created from A) EC-profiles and B) pathway-profiles. Note that in the pathway-profiles the difference is less significant.

**Table 1.** The meta-genomic/transcriptomic pathways that significantly correlate with the organic acids levels in the late spring samples.

| Pathway | Oxalate |  | Acetate |  | Formate* |  |
| --- | --- | --- | --- | --- | --- | --- |
|  | *r* | *p-value* | *r* | *p-value* | *r* | *p-value* |
| Fatty acid biosynthesis | 0.615 | 0.001 | 0.600 | 0.002 | 0.555 | 0.005 |
| Biosynthesis of unsaturated fatty acids | 0.736 | 0.000 | 0.647 | 0.001 | 0.488 | 0.015 |
| Fatty acid elongation | 0.622 | 0.001 | 0.529 | 0.008 | 0.348 | 0.096 |
| Geraniol degradation | 0.622 | 0.001 | 0.529 | 0.008 | 0.348 | 0.096 |
| Styrene degradation | -0.449 | 0.028 | -0.460 | 0.024 | -0.398 | 0.054 |

\*Formate significantly correlates with fatty acid elongation, geraniol degradation and styrene degradation at alpha level of 0.1.

The discrepancy in annotation between metagenomes (DNA) and metatranscriptomes (cDNA) has previously been observed in environments such as human gut [42] and open ocean [43]. The genes observed in the metagenomes represent potential functions that may or may not be expressed in the environment at the time of sampling and could belong to inactive community members; the metatranscriptome-specific functions belong to active members of the community at the time of sampling [44]. The low metagenome-to-transcriptome similarity in the early spring samples (Figure 2) suggest that the active members in early spring occur at such low abundance that metagenomic sequencing fails to detect them. We speculated that the potential functional diversity in the early spring metagenome samples (Figure 1; SOM Figure 1; DNA datasets) might come from the DNA of dead or inactive cells preserved in the snow. Interestingly, many microorganisms identified in snow and ice via 16S rRNA gene surveys are non-psychrophiles [45] and their importance in the community needs further investigation. Meanwhile, the diversity of active microbial functionality in the early spring metatranscriptomes (Figure 1; SOM Figure 1; RNA datasets) reflected diverse microbial activities (238 enzymatic functions involved in 84 metabolic pathways including cell size reduction, changes in fatty acid and phospholipid membrane composition, and decrease in the fractional volume of cellular water). This observation is in line with the known variety of survival strategies employed by microbes at low temperature [2, 17].

With the warming in the late spring, the active community made up a larger fraction of the sequenced reads and, thus, manifested in more homogeneity. Previous 16S rRNA-based taxonomic analysis on the same dataset observed a shift in the community from early to late spring [33]. While the early spring samples contained a core community of 59 OTUs, there were only 29 OTUs in the late spring samples, with 42 early spring core OTUs disappearing from the core community of late spring samples [33]. The early spring community contained a higher diversity of core organisms of which only a small fraction were likely active, and the inactive community members could no longer be detected in the late spring samples. As a result, we observed a decrease in the function diversity (Figure 1; SOM Figure 1) and an increase in the metagenome-to-transcriptome similarity (Figure 2). In addition, our result also suggests that despite the taxonomic diversity in the late spring samples, their functional capacity and activity were highly similar (Figure 1; SOM Figure 1; RNA datasets), highlighting the advantages of functional -omics analysis to the 16S rRNA gene surveys.

### 3.2. Microbial use of complex organic compounds in the snow

Snow provides a medium and nutrients for microbial growth and associated physicochemical processes [46], and growth implies the utilization of nutrients. Numerous genes detected in environmental ice metagenomes related to xenobiotics, biopolymers and other carbon sources suggest that glacial ice microorganisms have the potential to degrade a wide range of substrates [47]. Our hypothesis of increased active microbial community members in the late spring snow might be related to the changes in organic acid levels in the samples (oxalate, acetate and formate; SOM Table 2). All three organic acids remained low in concentration in the early spring samples. They increased in the late spring (SOM Figure 2), possibly concomitant with increased microbial activity. Microbial preferences for different carbon classes was studied in Antarctic snow and results showed a higher rate of carbon uptake when snow microcosms were amended with a combination of simple and complex carbon sources [48]. The appearance of organic acids in the snow may have both abiotic (e.g. aerial deposition) and biotic (e.g. microbial activity) origins. In our study, however, the clear correlation of their per-sample concentrations with different microbial activity levels, captured by metatranscriptomes, strongly indicates active metabolism in the late spring samples. Note that both EC-profiles and pathway-profiles correlated with the organic acid levels more significantly than the EggNog Mapper derived functional profiles (Methods; SOM Table 3) from Bergk-Pinto et al. [33], highlighting mi-faser’s adequacy for this study.

Among the enzymes that were not mapped to known KEGG pathways, two tRNA-methyltransferases (2.1.1.61 and 2.1.1.217; p-value<0.05, Methods) showed significant correlation with organic acid levels. tRNA methylation regulates important steps in protein synthesis and is essential for microbial growth in high temperature [49]. Our results suggest that it could be also involved in low temperature conditions.

We further identified five pathways in our meta-genomes/transcriptomes that significantly correlated with the organic acid levels in the late spring samples (Table 1; SOM Figure 3-7; p-value<0.05, Methods): fatty acid biosynthesis, biosynthesis of unsaturated fatty acids, fatty acid elongation, geraniol degradation, and styrene degradation. The top three pathways were related to fatty acid synthesis and elongation. Fatty acids are essential for living organisms due to their role in membrane synthesis, which is even more critical when low temperature affects membrane fluidity [50]. Geraniol degradation is important as geraniol is a terpene produced by a variety of plants for its antibacterial activities [51]. Terpenes are released from plants to the atmosphere [52], and deposited in arctic snowpacks like other volatile organic compounds [53]. Some bacteria, e.g. *Pseudomonas putida*, are able to utilize geraniol as their sole carbon and energy source [54]. P. putida is also known to degrade styrene [55] and polystyrene [56]. Therefore, the correlation and anticorrelation between the organic acids levels and, respectively, the microbial geraniol degradation and styrene degradation, may suggest a switch of nutrients in the environment. *P. putida* is known to possess diverse metabolic capabilities to degrade a variety of organic solvents. Most of its strains are mesophilic, but one (KT2440) has been reported as psychrotolerant (optimal growth at 30°C but can proliferate at 4°C) [57]. To the best of our knowledge, no microbial metabolism of geraniol and styrene has been reported at low temperatures with evidence at transcription level. Our functional –omics study thus indicates, for the first time, the activity of microbial degradation pathways of complex organic compounds at sub-zero temperatures.

## 4. Conclusions

We defined microbial activity at low temperature at the gene expression level in metagenomic and metatranscriptomic datasets from snow in early and late spring. Our results highlight the novel microbial activity of complex organic compound degradations at low temperature. Further in-depth exploration of the functionality of the cryosphere inhabitants can contribute to our understanding of microbial metabolism at low temperatures and aid in the discovery of novel enzymes with potential industrial and bioremediation value.

## Author Contributions

Conceptualization, Chengsheng Zhu and Yana Bromberg; Data curation, Benoit Bergk Pinto and Lorrie Maccario; Formal analysis, Chengsheng Zhu; Funding acquisition, Yana Bromberg; Resources, Yana Bromberg; Software, Maximilian Miller and Nicholas Lusskin; Supervision, Yana Bromberg; Visualization, Chengsheng Zhu; Writing – original draft, Chengsheng Zhu; Writing – review & editing, Chengsheng Zhu, Benoit Bergk Pinto, Max Haggblom, Timothy Vogel, Catherine Larose and Yana Bromberg.

## Acknowledgments

We thank Dr. Yannick Mahlich, Yanran Wang and Zishuo Zeng (Rutgers University) for the useful discussion and suggestions.

## References

1. Rothschild LJ, Mancinelli RL. Life in extreme environments. Nature. 2001;409:1092. doi: 10.1038/35059215.

2. Price PB, Sowers T. Temperature dependence of metabolic rates for microbial growth, maintenance, and survival. Proceedings of the National Academy of Sciences of the United States of America. 2004;101(13):4631–6. doi: 10.1073/pnas.0400522101. PubMed PMID: PMC384798.

3. Christner BC, Mosley-Thompson E, Thompson LG, Zagorodnov V, Sandman K, Reeve JN. Recovery and Identification of Viable Bacteria Immured in Glacial Ice. Icarus. 2000;144(2):479–85. doi: https://doi.org/10.1006/icar.1999.6288.

4. Christner BC, Mosley-Thompson E, Thompson LG, Reeve JN. Isolation of bacteria and 16S rDNAs from Lake Vostok accretion ice. Environmental Microbiology. 2001;3(9):570–7. doi: 10.1046/j.1462-2920.2001.00226.x.

5. Cameron KA, Stibal M, Zarsky JD, Gozdereliler E, Schostag M, Jacobsen CS. Supraglacial bacterial community structures vary across the Greenland ice sheet. FEMS microbiology ecology. 2016;92(2). Epub 2015/12/23. doi: 10.1093/femsec/fiv164. PubMed PMID: 26691594.

6. Uetake J, Tanaka S, Segawa T, Takeuchi N, Nagatsuka N, Motoyama H, et al. Microbial community variation in cryoconite granules on Qaanaaq Glacier, NW Greenland. FEMS microbiology ecology. 2016;92(9). Epub 2016/06/17. doi: 10.1093/femsec/fiw127. PubMed PMID: 27306554.

7. Webster-Brown JG, Hawes I, Jungblut AD, Wood SA, Christenson HK. The effects of entombment on water chemistry and bacterial assemblages in closed cryoconite holes on Antarctic glaciers. FEMS microbiology ecology. 2015;91(12). Epub 2015/11/18. doi: 10.1093/femsec/fiv144. PubMed PMID: 26572547.

8. Brinkmeyer R, Knittel K, Jurgens J, Weyland H, Amann R, Helmke E. Diversity and structure of bacterial communities in Arctic versus Antarctic pack ice. Applied and environmental microbiology. 2003;69(11):6610–9. Epub 2003/11/07. PubMed PMID: 14602620; PubMed Central PMCID: PMCPMC262250.

9. Amato P, Hennebelle R, Magand O, Sancelme M, Delort AM, Barbante C, et al. Bacterial characterization of the snow cover at Spitzberg, Svalbard. FEMS microbiology ecology. 2007;59(2):255–64. Epub 2007/03/03. doi: 10.1111/j.1574-6941.2006.00198.x. PubMed PMID: 17328766.

10. Larose C, Berger S, Ferrari C, Navarro E, Dommergue A, Schneider D, et al. Microbial sequences retrieved from environmental samples from seasonal arctic snow and meltwater from Svalbard, Norway. Extremophiles : life under extreme conditions. 2010;14(2):205–12. Epub 2010/01/13. doi: 10.1007/s00792-009-0299-2. PubMed PMID: 20066448.

11. Wunderlin T, Ferrari B, Power M. Global and local-scale variation in bacterial community structure of snow from the Swiss and Australian Alps. FEMS microbiology ecology. 2016;92(9). Epub 2016/06/15. doi: 10.1093/femsec/fiw132. PubMed PMID: 27297721.

12. Harding T, Jungblut AD, Lovejoy C, Vincent WF. Microbes in High Arctic Snow and Implications for the Cold Biosphere. Applied and environmental microbiology. 2011;77(10):3234. doi: 10.1128/AEM.02611-10.

13. Maccario L, Carpenter SD, Deming JW, Vogel TM, Larose C. Sources and selection of snow-specific microbial communities in a Greenlandic sea ice snow cover. Scientific Reports. 2019;9(1):2290. doi: 10.1038/s41598-019-38744-y.

14. Larose C, Prestat E, Cecillon S, Berger S, Malandain C, Lyon D, et al. Interactions between Snow Chemistry, Mercury Inputs and Microbial Population Dynamics in an Arctic Snowpack. PLOS ONE. 2013;8(11):e79972. doi: 10.1371/journal.pone.0079972.

15. Segawa T, Miyamoto K, Ushida K, Agata K, Okada N, Kohshima S. Seasonal change in bacterial flora and biomass in mountain snow from the Tateyama Mountains, Japan, analyzed by 16S rRNA gene sequencing and real-time PCR. Applied and environmental microbiology. 2005;71(1):123–30. Epub 2005/01/11. doi: 10.1128/aem.71.1.123-130.2005. PubMed PMID: 15640179; PubMed Central PMCID: PMCPMC544271.

16. Zhang S, Yang G, Wang Y, Hou S. Abundance and community of snow bacteria from three glaciers in the Tibetan Plateau. Journal of Environmental Sciences. 2010;22(9):1418–24. doi: https://doi.org/10.1016/S1001-0742(09)60269-2.

17. Nikrad MP, Kerkhof LJ, Haggblom MM. The subzero microbiome: microbial activity in frozen and thawing soils. FEMS microbiology ecology. 2016;92(6):fiw081. Epub 2016/04/24. doi: 10.1093/femsec/fiw081. PubMed PMID: 27106051.

18. Maccario L, Vogel TM, Larose C. Potential drivers of microbial community structure and function in Arctic spring snow. Frontiers in Microbiology. 2014;5:413. doi: 10.3389/fmicb.2014.00413. PubMed PMID: PMC4124603.

19. McNeil AJ. PROFILE: Early Excellence in Physical Organic Chemistry. Journal of Physical Organic Chemistry. 2012;25(8):611–. doi: 10.1002/poc.2969.

20. Ariya PA, Domine F, Kos G, Amyot M, Côté V, Vali H, et al. Snow – a photobiochemical exchange platform for volatile and semi-volatile organic compounds with the atmosphere. Environmental Chemistry. 2011;8(1):62–73. doi: https://doi.org/10.1071/EN10056.

21. Vogel TM, Simonet P, Jansson JK, Hirsch PR, Tiedje JM, van Elsas JD, et al. TerraGenome: a consortium for the sequencing of a soil metagenome. Nature Reviews Microbiology. 2009;7:252. doi: 10.1038/nrmicro2119.

22. Rusch DB, Halpern AL, Sutton G, Heidelberg KB, Williamson S, Yooseph S, et al. The Sorcerer II Global Ocean Sampling Expedition: Northwest Atlantic through Eastern Tropical Pacific. PLOS Biology. 2007;5(3):e77. doi: 10.1371/journal.pbio.0050077.

23. Human Microbiome Project Consortium. Structure, function and diversity of the healthy human microbiome. Nature. 2012;486(7402):207–14. Epub 2012/06/16. doi: 10.1038/nature11234. PubMed PMID: 22699609; PubMed Central PMCID: PMCPMC3564958.

24. Sunagawa S, Coelho LP, Chaffron S, Kultima JR, Labadie K, Salazar G, et al. Ocean plankton. Structure and function of the global ocean microbiome. Science (New York, NY). 2015;348(6237):1261359. Epub 2015/05/23. doi: 10.1126/science.1261359. PubMed PMID: 25999513.

25. Qin J, Li Y, Cai Z, Li S, Zhu J, Zhang F, et al. A metagenome-wide association study of gut microbiota in type 2 diabetes. Nature. 2012;490(7418):55–60. Epub 2012/10/02. doi: 10.1038/nature11450. PubMed PMID: 23023125.

26. Gevers D, Kugathasan S, Denson LA, Vazquez-Baeza Y, Van Treuren W, Ren B, et al. The treatment-naive microbiome in new-onset Crohn’s disease. Cell host & microbe. 2014;15(3):382–92. Epub 2014/03/19. doi: 10.1016/j.chom.2014.02.005. PubMed PMID: 24629344; PubMed Central PMCID: PMCPMC4059512.

27. Zhu C, Miller M, Marpaka S, Vaysberg P, Rühlemann MC, Wu G, et al. Functional sequencing read annotation for high precision microbiome analysis. Nucleic Acids Research. 2018;46(4):e23–e. doi: 10.1093/nar/gkx1209.

28. Hemme CL, Tu Q, Shi Z, Qin Y, Gao W, Deng Y, et al. Comparative metagenomics reveals impact of contaminants on groundwater microbiomes. Frontiers in Microbiology. 2015;6(1205). doi: 10.3389/fmicb.2015.01205.

29. Chen L-x, Hu M, Huang L-n, Hua Z-s, Kuang J-l, Li S-j, et al. Comparative metagenomic and metatranscriptomic analyses of microbial communities in acid mine drainage. The Isme Journal. 2014;9:1579. doi: 10.1038/ismej.2014.245 https://www.nature.com/articles/ismej2014245#supplementary-information.

30. Rodriguez-R LM, Overholt WA, Hagan C, Huettel M, Kostka JE, Konstantinidis KT. Microbial community successional patterns in beach sands impacted by the Deepwater Horizon oil spill. ISME J. 2015;9(9):1928–40. doi: 10.1038/ismej.2015.5.

31. Zhu C, Delmont TO, Vogel TM, Bromberg Y. Functional Basis of Microorganism Classification. PLOS Computational Biology. 2015;11(8):e1004472. doi: 10.1371/journal.pcbi.1004472.

32. Zhu C, Mahlich Y, Miller M, Bromberg Y. fusionDB: assessing microbial diversity and environmental preferences via functional similarity networks. Nucleic Acids Research. 2018;46(D1):D535–D41. doi: 10.1093/nar/gkx1060.

33. Bergk Pinto B, Maccario L, Dommergue A, Vogel TM, Larose C. Do Organic Substrates Drive Microbial Community Interactions in Arctic Snow? Frontiers in Microbiology. 2019;10(2492). doi: 10.3389/fmicb.2019.02492.

34. Schloss PD, Westcott SL, Ryabin T, Hall JR, Hartmann M, Hollister EB, et al. Introducing mothur: Open-Source, Platform-Independent, Community-Supported Software for Describing and Comparing Microbial Communities. Applied and environmental microbiology. 2009;75(23):7537. doi: 10.1128/AEM.01541-09.

35. Schloss PD, Gevers D, Westcott SL. Reducing the Effects of PCR Amplification and Sequencing Artifacts on 16S rRNA-Based Studies. PLOS ONE. 2011;6(12):e27310. doi: 10.1371/journal.pone.0027310.

36. Andrews S. FastQC: a quality control tool for high throughput sequence data. 2010.

37. Edgar RC. Search and clustering orders of magnitude faster than BLAST. Bioinformatics (Oxford, England). 2010;26(19):2460–1. Epub 2010/08/17. doi: 10.1093/bioinformatics/btq461. PubMed PMID: 20709691.

38. Miller M, Zhu C, Bromberg Y. clubber: removing the bioinformatics bottleneck in big data analyses. Journal of integrative bioinformatics. 2017;14(2). Epub 2017/06/14. doi: 10.1515/jib-2017-0020. PubMed PMID: 28609295.

39. Ec W. Enzyme nomenclature 1992: recommendations of the Nomenclature Committee of the International Union of Biochemistry and Molecular Biology on the nomenclature and classification of enzymes: Academic Press, San Diego, California; 1992.

40. Kanehisa M, Sato Y, Kawashima M, Furumichi M, Tanabe M. KEGG as a reference resource for gene and protein annotation. Nucleic Acids Research. 2016;44(Database issue):D457–D62. doi: 10.1093/nar/gkv1070. PubMed PMID: PMC4702792.

41. Best DJ, Roberts DE. Algorithm AS 89: The Upper Tail Probabilities of Spearman’s Rho. Applied Statistics. 1975;24:377–9.

42. Franzosa EA, Morgan XC, Segata N, Waldron L, Reyes J, Earl AM, et al. Relating the metatranscriptome and metagenome of the human gut. Proceedings of the National Academy of Sciences. 2014;111(22):E2329. doi: 10.1073/pnas.1319284111.

43. Shi Y, Tyson GW, Eppley JM, DeLong EF. Integrated metatranscriptomic and metagenomic analyses of stratified microbial assemblages in the open ocean. The Isme Journal. 2010;5:999. doi: 10.1038/ismej.2010.189 https://www.nature.com/articles/ismej2010189#supplementary-information.

44. Yu K, Zhang T. Metagenomic and Metatranscriptomic Analysis of Microbial Community Structure and Gene Expression of Activated Sludge. PLOS ONE. 2012;7(5):e38183. doi: 10.1371/journal.pone.0038183.

45. Cowan DA, Tow LA. Endangered Antarctic Environments. Annual Review of Microbiology. 2004;58(1):649–90. doi: 10.1146/annurev.micro.57.030502.090811. PubMed PMID: 15487951.

46. Domine F, Shepson PB. Air-snow interactions and atmospheric chemistry. Science (New York, NY). 2002;297(5586):1506–10. Epub 2002/08/31. doi: 10.1126/science.1074610. PubMed PMID: 12202818.

47. Stibal M, Šabacká M, Žárský J. Biological processes on glacier and ice sheet surfaces. Nature Geoscience. 2012;5:771. doi: 10.1038/ngeo1611.

48. Antony R, Krishnan KP, Laluraj CM, Thamban M, Dhakephalkar PK, Engineer AS, et al. Diversity and physiology of culturable bacteria associated with a coastal Antarctic ice core. Microbiol Res. 2012;167(6):372–80. Epub 2012/04/28. doi: 10.1016/j.micres.2012.03.003. PubMed PMID: 22537873.

49. Hori H. Methylated nucleosides in tRNA and tRNA methyltransferases. Frontiers in Genetics. 2014;5:144. doi: 10.3389/fgene.2014.00144. PubMed PMID: PMC4033218.

50. Cronan JE, Thomas J. Bacterial Fatty Acid Synthesis and its Relationships with Polyketide Synthetic Pathways. Methods in enzymology. 2009;459:395–433. doi: 10.1016/S0076-6879(09)04617-5. PubMed PMID: PMC4095770.

51. Friedman M, Henika PR, Mandrell RE. Bactericidal activities of plant essential oils and some of their isolated constituents against Campylobacter jejuni, Escherichia coli, Listeria monocytogenes, and Salmonella enterica. Journal of food protection. 2002;65(10):1545–60. Epub 2002/10/17. PubMed PMID: 12380738.

52. Marmulla R, Harder J. Microbial monoterpene transformations-a review. Front Microbiol. 2014;5:346. Epub 2014/08/01. doi: 10.3389/fmicb.2014.00346. PubMed PMID: 25076942; PubMed Central PMCID: PMCPMC4097962.

53. Kos G, Kanthasami V, Adechina N, Ariya PA. Volatile organic compounds in Arctic snow: concentrations and implications for atmospheric processes. Environmental science Processes & impacts. 2014;16(11):2592–603. Epub 2014/09/25. doi: 10.1039/c4em00410h. PubMed PMID: 25249335.

54. Vandenbergh PA, Wright AM. Plasmid Involvement in Acyclic Isoprenoid Metabolism by Pseudomonas putida. Applied and environmental microbiology. 1983;45(6):1953–5. Epub 1983/06/01. PubMed PMID: 16346325; PubMed Central PMCID: PMCPMC242567.

55. O’Connor K, Duetz W, Wind B, Dobson AD. The effect of nutrient limitation on styrene metabolism in Pseudomonas putida CA-3. Applied and environmental microbiology. 1996;62(10):3594–9. Epub 1996/10/01. PubMed PMID: 8967774; PubMed Central PMCID: PMCPMC168165.

56. Ward PG, Goff M, Donner M, Kaminsky W, O’Connor KE. A Two Step Chemo-biotechnological Conversion of Polystyrene to a Biodegradable Thermoplastic. Environmental Science & Technology. 2006;40(7):2433–7. doi: 10.1021/es0517668.

57. Fonseca P, Moreno R, Rojo F. Growth of Pseudomonas putida at low temperature: global transcriptomic and proteomic analyses. Environmental microbiology reports. 2011;3(3):329–39. Epub 2011/06/01. doi: 10.1111/j.1758-2229.2010.00229.x. PubMed PMID: 23761279.

